## Supplementary Materials for "The COMBO window: A chronic cranial implant for multiscale circuit interrogation in mice"

#### Files:

**Supplementary File 1:** COMBO_cup.stl
**Supplementary File 2:** COMBO_flat.stl
**Supplementary File 3:** COMBO_lateral.stl
**Supplementary File 4:** COMBO_posterior.stl
**Supplementary File 5:** COMBO_anterior.stl
**Supplementary File 6:** COMBO_Q1.stl
**Supplementary File 7:** COMBO_Q2.stl
**Supplementary File 8:** COMBO_Q3.stl
**Supplementary File 9:** COMBO_Q4.stl
**Supplementary File 10:** Head_plate.dwg
**Supplementary File 11:** Head_plate_holder.sldprt
**Supplementary File 12:** Head_plate_holder_top.dwg
**Supplementary File 13:** Head_plate_holder_bottom.dwg
**Supplementary File 14:** Brain_mold.stl

#### Videos:

**Supplementary Video 1:** Facial videography during a trial of sucrose delivery
**Supplementary Video 2:** Facial videography during a trial of quinine delivery
**Supplementary Video 3:** Brain-wide fUS activity in awake mice in response to visual stimulation
**Supplementary Video 4:** Facial videography and two-photon imaging in the retrosplenial cortex during locomotion

#### Figures:

**Supplementary Figure 1:** Head fixation part details
**Supplementary Figure 2:** COMBO window assembly and installation instructions
**Supplementary Figure 3:** Sporadic GFAP fluorescence was observed in mice implanted with the COMBO window
**Supplementary Figure 4:** Behavioral effects are consistent across sex and individuals
**Supplementary Figure 5:** The COMBO window enables chronic brain-wide acquisition of fUS data
**Supplementary Figure 6:** The COMBO window facilitates longitudinal experiments to interrogate neural circuits underlying behavior.

#### Tables:

**Supplementary Table 1**: Open field foraging task statistical analysis
**Supplementary Table 2**: Facial expression statistical analysis
**Supplementary Table 3**: Author Contributions

#### Appendix:

**Appendix 1**: COMBO window preparation and installation protocol

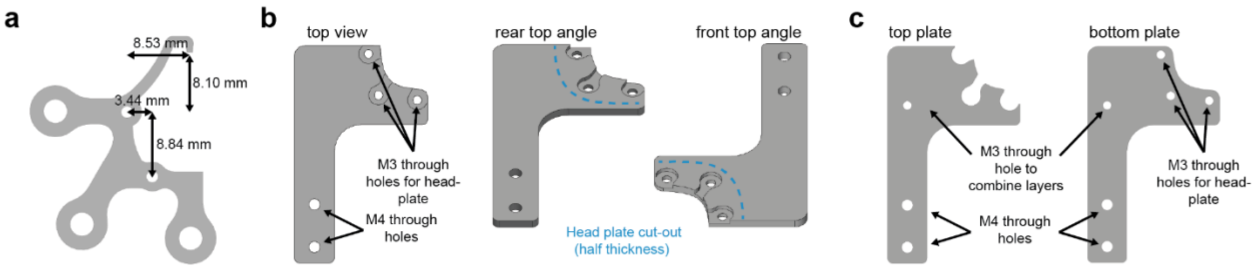

**Supplementary Figure 1: Head fixation part details**

**a** Computer-aided design of the standard COMBO window head-plate (Supplementary File 10). The head-plate attaches to the implant via two M1.4 through holes at the side and rear, as well as a peg in the front. Other custom head plate designs with the same features and relative distances can also be used for head fixation. **b** The standard head-plate holder design (Supplementary File 11) consists of a single metal plate with the head plate outline cut halfway through the total thickness. Threaded M3 screws are welded into the head-plate holes and grinded flush with the underside of the plate. M4 through holes allow for attachment to other commercial or custom parts for further stabilization. It is recommended that a machine shop helps with the fabrication of this part. **c** An alternative head-plate holder design consists of a top and bottom plate (Supplementary Files 12-13) that can each be laser cut and joined together with no custom fabrication. M3 screws can be used to secure the two layers together, and M4 screws to attach the head-plate holder to other commercial or custom parts. Additional M3 screws can be attached via the underside of the holder using glue/epoxy.

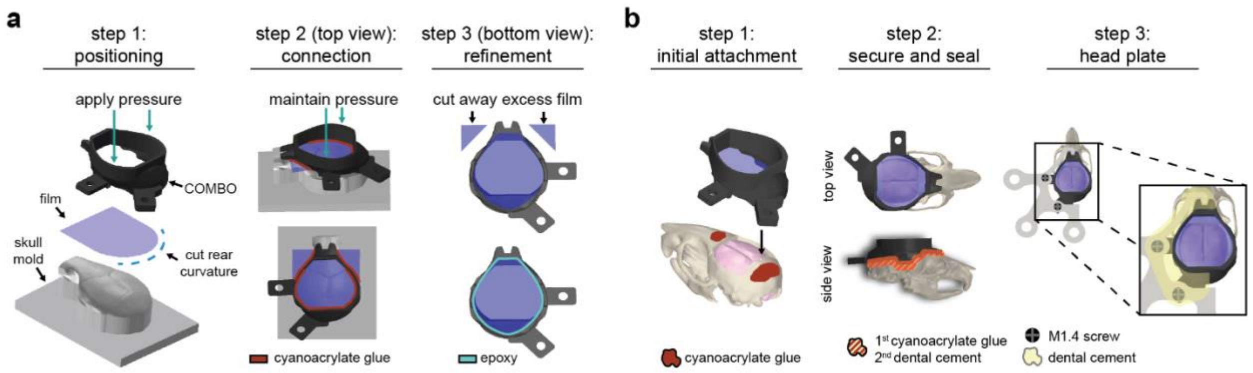

**Supplementary Figure 2: COMBO window assembly and installation instructions**

**a** Three-step diagram of the preparation of the COMBO window. Using the skull mold (Supplementary File 14) is recommended but not required for proper assembly. **b** Three-step diagram of the installation of the COMBO window after a cranial window has been created. The head plate can be installed at the same time as Steps 1-2 or at a later date. Detailed methods for both of the procedures are provided in Appendix 1.

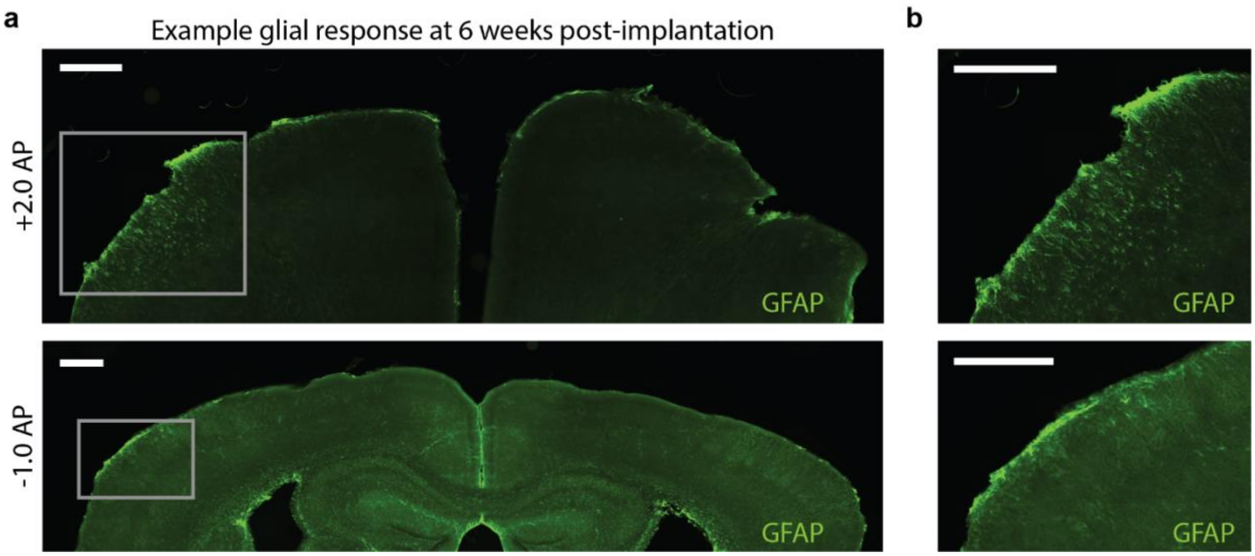

**Supplementary Figure 3: Sporadic GFAP fluorescence was observed in mice implanted with the COMBO window**

**a** Glial fibrillary acidic protein (GFAP) fluorescence in two example slices (top: bregma +2.0 mm AP, bottom: bregma -1.0 mm AP) of mice at 6 weeks after being implanted with the COMBO window. In both images, a localized increase of GFAP fluorescence can be seen in the left hemisphere. **b** Zoomed-in images of the elevated GFAP signal indicate that the immune response was found mostly in fibers located at or near the pial surface. Scale bars represent 500 µm.

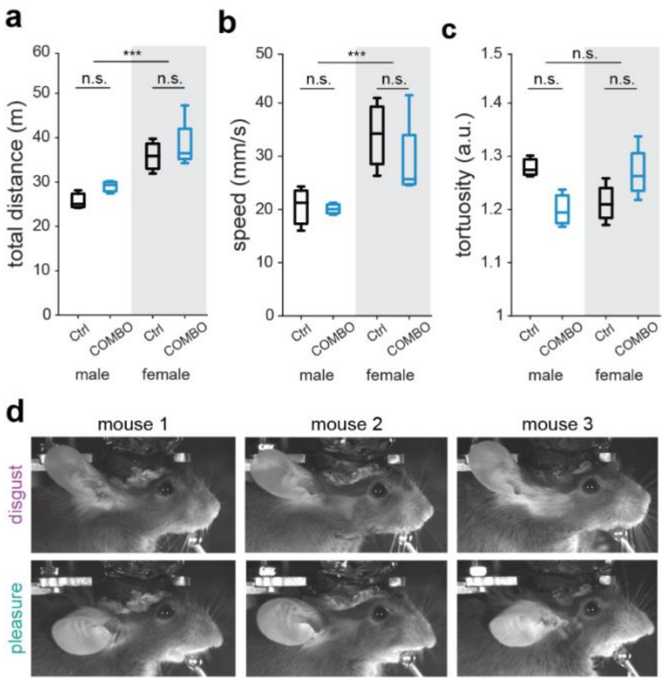

**Supplementary Figure 4: Behavioral effects are consistent across sex and individuals**

**a-c** The total distance (**a**), speed (**b**), and tortuosity (**c**) of control and COMBO window mice separated into male (n = 3) and females (n = 4). Boxplots represent the median (center line), 25th and 75th percentiles (lower and upper box), and the 1st and 99th percentile (whiskers). Two-way ANOVA on ranks with main effects of sex and cranial window. Main effect of sex: *** $p < 0.001$. Post hoc pairwise t-tests, Bonferroni corrected: n.s. $p > 0.05$. **d** Example prototypical disgust and pleasure facial expressions exhibited by three animals with a “cup” version of the COMBO window installed. Key features of the elicited disgust face include a flaring back of ear and an upturned snout, and of the elicited pleasure face include the forward movement of the ear and a downturned snout.

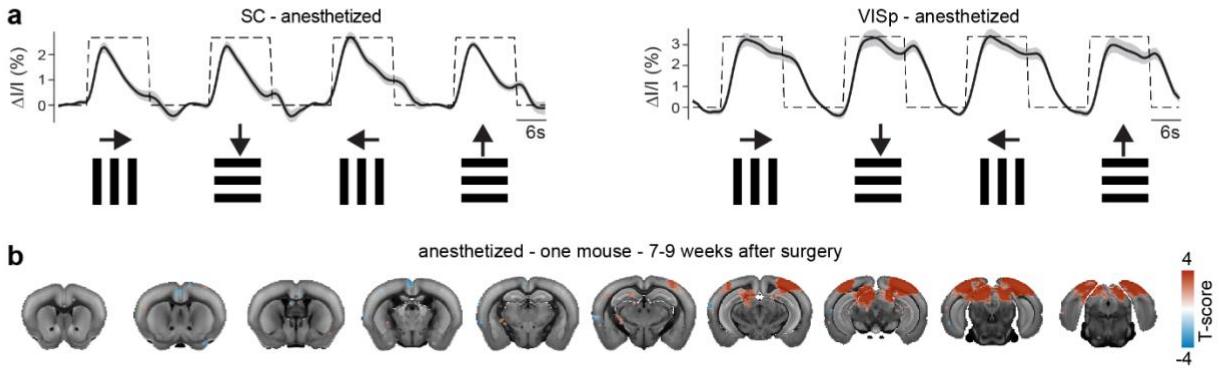

**Supplementary Figure 5: The COMBO window enables chronic brain-wide acquisition of fUS data**

**a** fUS signal in the superior colliculi (SC) and primary visual cortex (VIS) of anesthetized mice covaries in response to drifting gratings in all four cardinal directions (N = 5 mice, n = 32 sessions). The dark black line and light gray shaded area represent the mean ± s.e.m. across sessions. **b** GLM results from 8 sessions of a single mouse recorded 7-9 weeks after surgery overlayed on the Allen Brain Atlas. Only voxels with an average T-score > 2 are displayed.

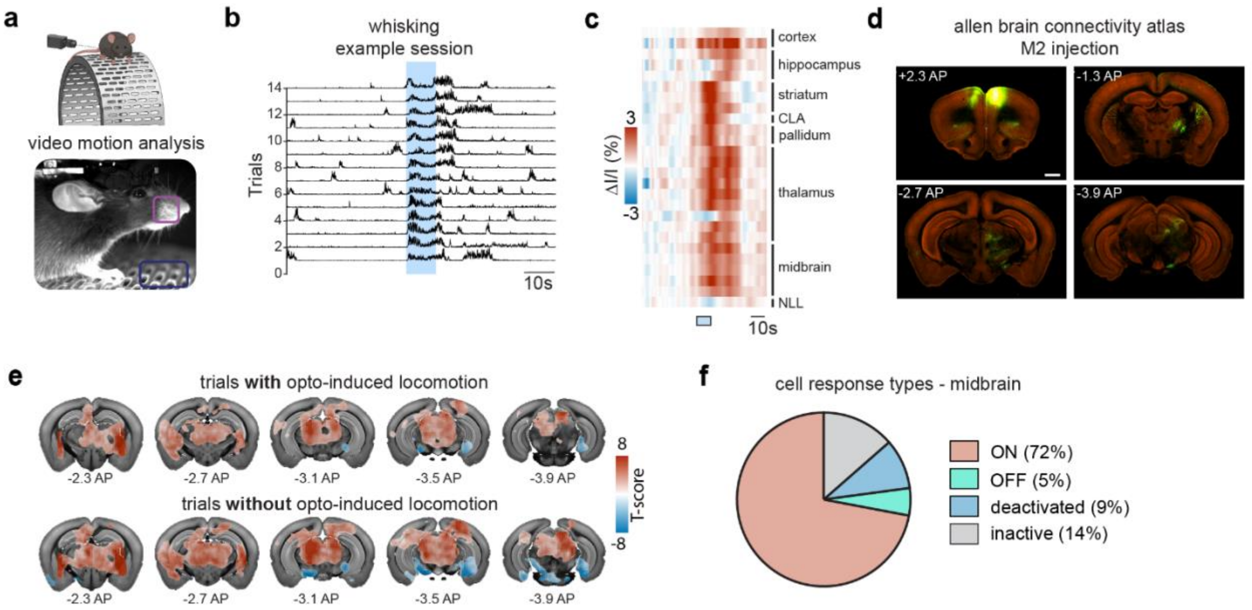

**Supplementary Figure 6: The COMBO window facilitates longitudinal experiments to interrogate neural circuits underlying behavior**

**a** Facial videography was used to monitor animal behavior on a running wheel. Regions-of-interest (ROIs) placed over the whisker pad (violet) and wheel (green) were utilized to capture whisking activity and locomotion, respectively, induced by optogenetic stimulation of the secondary motor cortex (M2). **b** Consecutive trials from the same example session as in Figure 5c showing a robust and reliable increase in whisking in response to optogenetic activation of M2. Each trial was z-scored to a pre-stimulus baseline and rescaled between 0 and 1 for visualization purposes. **c** Region-wise segmented results of the optogenetically-induced fUS activity. Regions are sorted by brain area, and only significantly modulated (correlation between stimulus timing and fUS signal) regions are included (significantly different from zero across sessions, $p < 0.001$, FDR-corrected). **d** Example coronal slices from the Allen Brain Connectivity Atlas (connectivity.brain-map.org/projection/experiment/287995889). AAV tracings after injection into the M2 (1) show widespread axonal projections from M2 to the striatum, the thalamus and the midbrain (2 - 4). Scale bar represents 1 mm. **e** GLM analysis of fUS data in response to optogenetic stimulation of M2 (N = 2 mice, n = 12 sessions). In contrast to Figure 5e, here the trials were separated according to strong or weak locomotor response (see methods for threshold definition) to optogenetic stimulation. Only the voxels with T-scores significantly different from zero across sessions ($p < 0.05$, FDR-corrected) are shown. **f** Pie chart showing the proportion of different cell response types observed in the midbrain (see methods for cell response type definition).

**Supplementary Table 1:** Open field foraging task statistical analysis

| Parameter | Statistical Test | Comparison | Post Hoc Test | P-value |
|---|---|---|---|---|
| Total Distance | Two-way ANOVA on ranks<br><br>CW vs Ctrl<br>F(1,13) = 1.93, p = 0.19<br><br>Male vs Female<br>F(1,13) = 34.05, ***p = 0.002 | COMBO vs Ctrl<br>(Male only) | Bonferroni | p = 1.00 |
| | | COMBO vs Ctrl<br>(Female only) | Bonferroni | p = 1.00 |
| Speed | Two-way ANOVA on ranks<br><br>CW vs Ctrl<br>F(1,13) = 0.75, p = 0.41<br><br>Male vs Female<br>F(1,13) = 31.21, ***p = 0.002 | COMBO vs Ctrl<br>(Male only) | Bonferroni | p = 1.00 |
| | | COMBO vs Ctrl<br>(Female only) | Bonferroni | p = 1.00 |
| Tortuosity | Two-way ANOVA on ranks<br><br>CW vs Ctrl<br>F(1,13) = 1.27, p = 0.29<br><br>Male vs Female<br>F(1,13) = 0.03, p = 0.86 | COMBO vs Ctrl<br>(Male only) | Bonferroni | p = 0.06 |
| | | COMBO vs Ctrl<br>(Female only) | Bonferroni | p = 0.48 |

**Supplementary Table 2:** Facial expression statistical analysis

| Parameter | Statistical Test | Comparison | P-value |
|---|---|---|---|
| Disgust | Wilcoxon rank sum test | Quinine vs Neutral | p = 0.0001 |
| | | Sucrose vs Neutral | p = 1.00 |
| Pleasure | Wilcoxon rank sum test | Quinine vs Neutral | p = 0.92 |
| | | Sucrose vs Neutral | p = 0.0011 |

**Supplementary Table 3:** Author Contributions

| | BJE | DS | PW | BJ | BS | AR | NG | TF | EM |
|---|---|---|---|---|---|---|---|---|---|
| **Conceptualization** | X | X | | | | | | | X |
| **Data Curation** | X | X | X | | | | | | |
| **Formal Analysis** | X | X | | | | | | | |
| **Funding Acquisition** | X | X | | | | | X | X | X |
| **Investigation** | X | X | X | X | X | X | | | |
| **Methodology** | X | X | X | X | | | X | X | X |
| **Project Administration** | X | | | | | | | | X |
| **Resources** | | | | | | | X | X | X |
| **Software** | X | X | | | | | | | X |
| **Supervision** | X | | | | | | X | X | X |
| **Validation** | X | X | X | | | | | | X |
| **Visualization** | X | X | | | | | | | X |
| **Writing - Original Draft** | X | X | | | | | | | X |
| **Writing - Review & Editing** | X | X | X | | | | | X | X |

### Appendix 1: COMBO window preparation and installation protocol

#### 1. COMBO Window Preparation

1.1. Download and print one of the COMBO window files (**Supplementary Files S1-9**) and the Brain_mold.stl (**Supplementary File 14)** using a 3D printer.
